# Dopamine transporter is a master regulator of dopaminergic neural network connectivity

**DOI:** 10.1101/2021.01.22.427804

**Authors:** Douglas Miller, Dylan T. Guenther, Andrew P. Maurer, Carissa A. Hansen, Andrew Zalesky, Habibeh Khoshbouei

## Abstract

Dopaminergic neurons of the substantia nigra (SNC) and ventral tegmental area (VTA) exhibit spontaneous firing activity. The dopaminergic neurons in these regions have been shown to exhibit differential sensitivity to neuronal loss and psychostimulants targeting dopamine transporter. However, it remains unclear whether these regional differences scale beyond individual neuronal activity to regional neuronal networks. Here we utilized live-cell calcium imaging to show that network connectivity greatly differs between SNC and VTA regions with higher incidence of hub-like neurons in the VTA. Specifically, the frequency of hub-like neurons was significantly lower in SNC dopamine neurons than in the adjacent VTA, consistent with the interpretation of a lower network resilience to SNC neuronal loss. We tested this hypothesis when activity of an individual dopaminergic neuron is suppressed, through whole-cell patch clamp electrophysiology, in either SNC, or VTA networks. Neuronal loss in the SNC decreased network clustering, whereas the larger number of hub-neurons in the VTA overcompensated by increasing network clustering in the VTA. We further show that network properties are regulatable via a dopamine transporter but not a D2 receptor dependent mechanism. Our results demonstrate novel regulatory mechanisms of functional network topology in dopaminergic brain regions.

## Introduction

Dopaminergic neurons are biochemically^1,2^, structurally^3,4^, and functionally heterogeneous^5–7^ and involved in a constellation of behaviors such as movement and value-based choice^8–11^. Transcriptomic and molecular studies have identified multiple distinct classes of dopaminergic neurons that exist within the same anatomical region^121,2,5,13–27^. Morphological investigations revealed a diverse and complex variation in the structural compartmentalization of dopamine neurons within each anatomical cluster.

The substantia nigra (SN, A9) and adjacent ventral tegmental area (VTA, A10) harbor two major dopaminergic clusters^3,4^. Despite their proximity, these two ventral midbrain regions contain strikingly different biological attributes, dopaminergic morphologies and protein expression patterns^1–4^. While these differences have been hypothesized to underlie differential sensitivity to neurotoxins^28–31^, drugs of abuse^4,6,32,33^, and neurodegeneration^34–38^, it remains unclear how these differences manifest within the VTA and SN regional networks.

Furthermore, dopaminergic neurons vary in their intrinsic electrophysiological properties^6,39,40^, such as spontaneous firing rate, excitability, and spike duration, suggesting that individual neurons may differentially contribute to the synchronization of activity in dopaminergic networks. However, it remains unknown how networks arise from local interactions of dopaminergic neurons, what properties these regional neuronal networks exhibit, whether they differ by specific midbrain regional (VTA vs. SNC) identity, how they are regulated or fluctuate in time.

Recently, network neuroscience has emerged as an important tool to answer integral questions about brain network organization^41^. Previous work utilizing network theory found that densely connected neurons (hubs) not only exist but orchestrate synchrony^42–45^. Recent works have suggested that dopamine network synchrony drives interregional connectivity and whole brain synchronization^8,46^. However, dopaminergic networks have yet to be investigated for the relative impact of the activity of individual neurons to network topology. Hubs, or highly connected neurons, are a common feature of complex networks^42,43,47–49^, but dopaminergic networks have yet to be tested empirically for the existence of hub neurons. Consistently, many investigations describe the substantia nigra pars compacta dopaminergic network as more vulnerable than the adjacent ventral tegmental area, albeit with a less understood mechanism^30,50–57^. While the origin of these regional differences has remained nebulous, network analysis can provide clues to how functional network organization of these regions may contribute to resilience or failure within the network that ultimately lead to neurocognitive and behavioral dysfunction.

The dopamine transporter, the spatial and temporal regulator of dopamine signaling^58–64^, which is expressed in nearly every dopamine neuron in both ventral tegmental area and substantia nigra, providing a unique modicum of communication that is specific to monoamine systems. It has been shown previously that stimulation of dopamine transporter induces alteration in local network properties in reduced systems^65^, and repeated exposure to modifiers of dopaminergic signaling, such as psychostimulants, regulates dopaminergic signaling machinery and brain activity^65–74^. Dopaminergic signaling relies heavily on volume transmission^64,75–78^, enabling communication between both adjacent and distant neurons^77,79–82^. Termination of the dopaminergic signal is thereby achieved by reuptake through dopamine transporter. Yet, it remains unexplored how dopamine transporter activity regulates interactions between dopaminergic neurons, leading to overall regulation of functional connectivity and ultimately dopaminergic network properties within the SNC and VTA. Previous models of dopaminergic networks suggest that information is encoded through either firing rate or synchronized activity^8,83,84^. However, these models do not explain if dopamine transporter function is critical to the functional structure of dopaminergic networks, or whether perturbations of dopamine transporter activity result in restructuring of the network.

In this work, we utilized high speed live-cell calcium imaging to investigate dynamic functional connectivity and network properties of the substantia nigra pars compacta and ventral tegmental area. Moreover, we determined regional-specific contributions of individual neurons to dopaminergic networks through electrophysiological manipulations and identified the role of dopamine transporter to the regulation of dopaminergic networks. We found that dopaminergic neurons in both VTA and SNC exhibit dynamic activity with similar magnitudes and variance of functional connectivity, but the incidence of hub-like neurons was significantly lower in substantia nigra dopamine neurons than in the adjacent ventral tegmental area. Consequently, suppression of activity of an individual dopaminergic neuron within the SNC network increased network clustering, which measures triplet interactions important for functional integration, whereas it decreased network clustering in the VTA. Furthermore, we found that methamphetamine stimulation of dopamine transporter alters network assortativity and small-world properties, which can contribute to network vulnerability to insults like aberrant protein aggregation or neurotoxins. These results suggest that the regional heterogeneities of dopaminergic networks influence functional network topology. Furthermore, differences in functional network topology contribute or ameliorate in the presence of neuronal loss, and the controllability of these networks through dopamine transporter regulation. Importantly, this work establishes a foundational exploration of the functional properties of dopaminergic network structure in these two dopaminergic nuclei and reveals how differences between regions may facilitate or reduce network dysfunction.

## Results

### Establishing a temporally and spatially-resolved model system to investigate the functional connectivity and network properties of dopaminergic neurons in the VTA and SNC

To investigate the intrinsic modulation of dynamic functional connectivity and properties of dopaminergic networks of the ventral tegmental area (VTA) and substantia nigra pars compacta (SNC), we utilized highspeed calcium imaging of neural activity (Fig 1). Dopaminergic neurons signal through their canonical neurotransmitter, dopamine, at multiple release sites, notably through somatodendritic release via a calcium-dependent manner^64,85^. Furthermore, dopaminergic signalling occurs at multiple timescales^9,86^, self-signaling^59,80,82,87–90^, and mechanisms outside of receptor-dependent modulation of activity^59,62,91^. While dopaminergic neurons receive diverse inputs into their respective regions, little is known about the self-organized intrinsic dopaminergic networks of the VTA and SNC. For these reasons, we utilized live-cell calcium imaging in acute midbrain slices spanning the VTA and SNC of mice, which provides a readout of neural activity that is intrinsically related to dopamine neuronal activity^92^. We achieved high-specificity expression of GCaMP6f in the dopamine neurons, within these regions, by crossing a GCaMP6f reporter line with DAT-cre. The recordings were performed at 20-25 Hz with an average of 13 individually resolvable neurons per experiment. Thereby, our model system provides a rigorous approach to resolve dopamine neuronal activity with both high spatial and temporal resolution afforded by GCaMP6f kinetics, as well as recording from both SNC and VTA within the same slice.

**Fig. 1:**
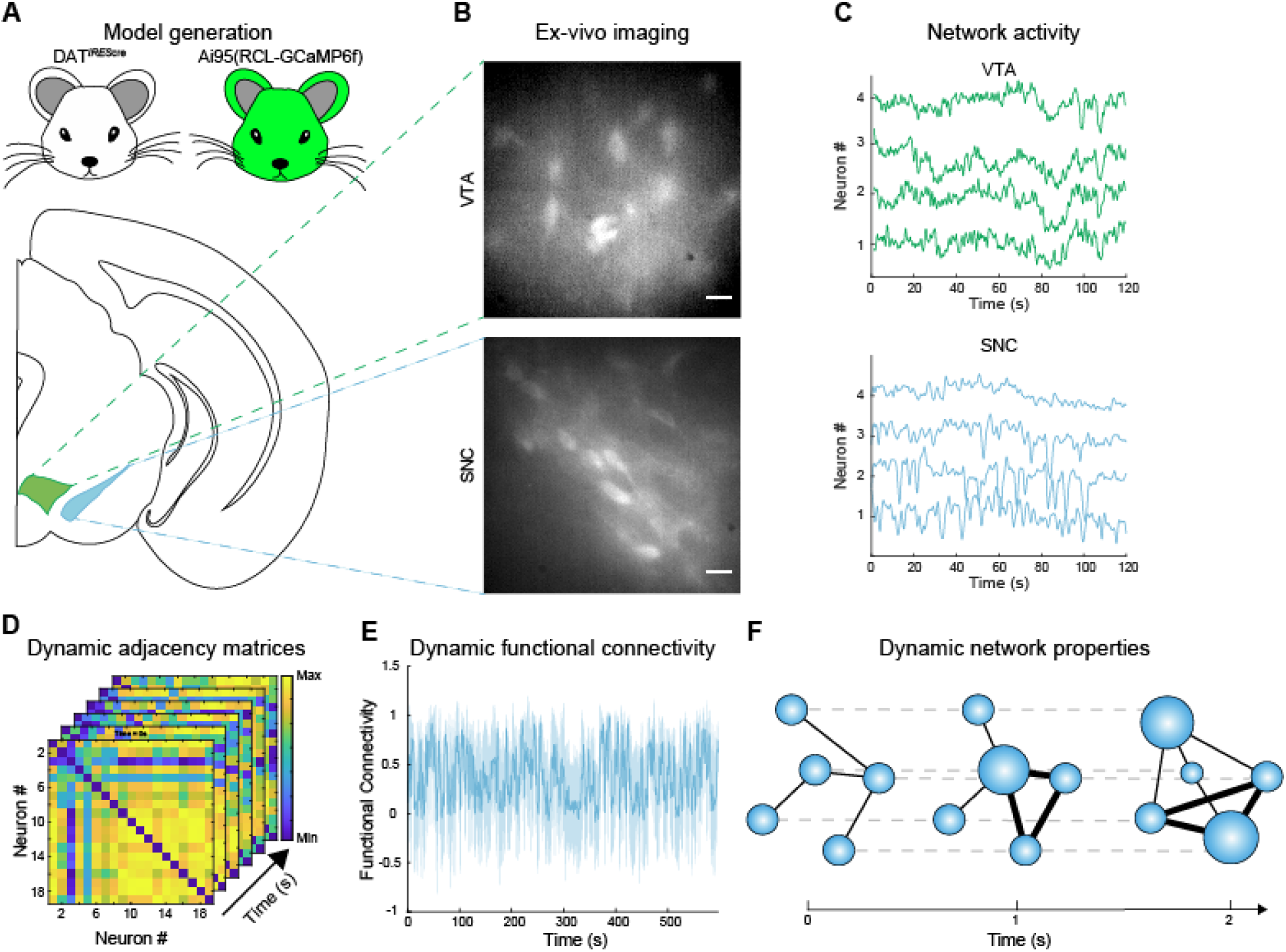
Schematic of experimental design and analytical pipeline to assay dynamic network properties in the VTA and SNC. Overview of experimental design and analytical processing. (**A**) DAT^*IREScre*^ mice were crossed with Ai95(RCL-GCaMP6f) mice to yield mice expressing GCaMP6f in dopaminergic neurons of the ventral midbrain. (**B**) Imaging of spontaneous dopaminergic activity is performed in the substantia nigra pars compacta (SNC) and ventral tegmental area (VTA). Scale bar = 50 μm. (**C**) Visibly active neurons were manually segmented as regions of interest and spontaneous neuronal calcium signal activity was extracted. (**D**) Pairwise correlation coefficients (Spearman’s rho) were calculated between each neuron of a sliding temporal window (100 frames, 1 frame step size) to generate dynamic adjacency matrix stack. (**E**) Per-matrix averages were used to determine average functional connectivity over time. (**F**) Weighted networks, in which correlations between neurons are factored into network measurements, are constructed from dynamic adjacency matrices (D). To balance specificity (correlations higher than a percentile) and sensitivity (inclusion of weak correlations), network measurements were calculated over multiple thresholds, whereby 5% of the strongest correlations are retained for measurement. Thresholding proceeds in 1% increments until all correlations are retained (100% threshold). Final measurements are then averaged across all thresholds to produce the mean network topology per time point. Circles represent individual neurons and their size corresponds to node strength in the network topology (larger = more contributing). Connection thickness indicates correlation magnitude between neurons.

Calcium influx and GCaMP6f responses exhibit a nonlinear relationship^93^. Additionally, dopaminergic signalling occurs at multiple timescales, and synchronized neuronal activity is pivotal in dopamine communication^8,9,46,86^. For these reasons, we utilized Spearman’s rho on the calcium signal between every neuron within the field of view over a 100-frame sliding window to construct a stack of adjacency matrices. Each matrix of correlation coefficients is then averaged to give the instantaneous functional connectivity of the network.

The time resolved adjacency matrices were then used to construct dynamic networks. Network measurements have previously been shown to be particularly sensitive to threshold values^94,95^, e.g. correlations that are believed to be meaningful. Importantly, there is no consensus on a meaningful correlation^95,96^, i.e. a correlation of 0.2 could be as important as a value of 0.9; therefore, we strategized with a sliding threshold function where relative thresholds were applied starting at a retainer of 5% of the strongest correlations, to a fully connected network with all correlations retained, at 1% intervals.

At each level, network metrics were calculated and averaged where necessary to give a time-resolved network metric before moving to the next matrix. This procedure enabled detailed and comprehensive investigation of the dynamic properties of dopaminergic networks of the VTA and SNC, and how they are mechanistically regulated.

### Substantia nigra pars compacta and ventral tegmental area exhibit similar patterns of dynamic functional connectivity

Dynamic functional connectivity quantifies the coupling between the time-series activity of a pair of neurons within a network as a function of time. Traditionally, these are applied to time-locked behavior events^97^. However, dopaminergic neurons exhibit tonic spontaneous firing activity that perseveres when excitatory neurotransmission is blocked^62,98,99^, suggesting a basal pattern of activity occurs in each region. Here, we utilize correlated calcium signals to determine dynamic functional connectivity of SNC and VTA dopaminergic networks. First, we assayed the average functional connectivity (average of correlations between all neuronal pairs within the observed network) over the entire duration of the experiment within each region followed by analysis of the variance of this measurement across time to determine whether SNC and VTA exhibit similar functional connectivity over time. Surprisingly, the SNC and VTA exhibited similar average functional connectivity (Figure 2B, two-tailed unpaired t-test, p = 0.7139, n = 5-8 sections from 5 animals), suggesting that functional connectivity alone does not sufficiently provide an understanding of differential vulnerability between VTA and SNC dopaminergic neurons.

**Fig. 2:**
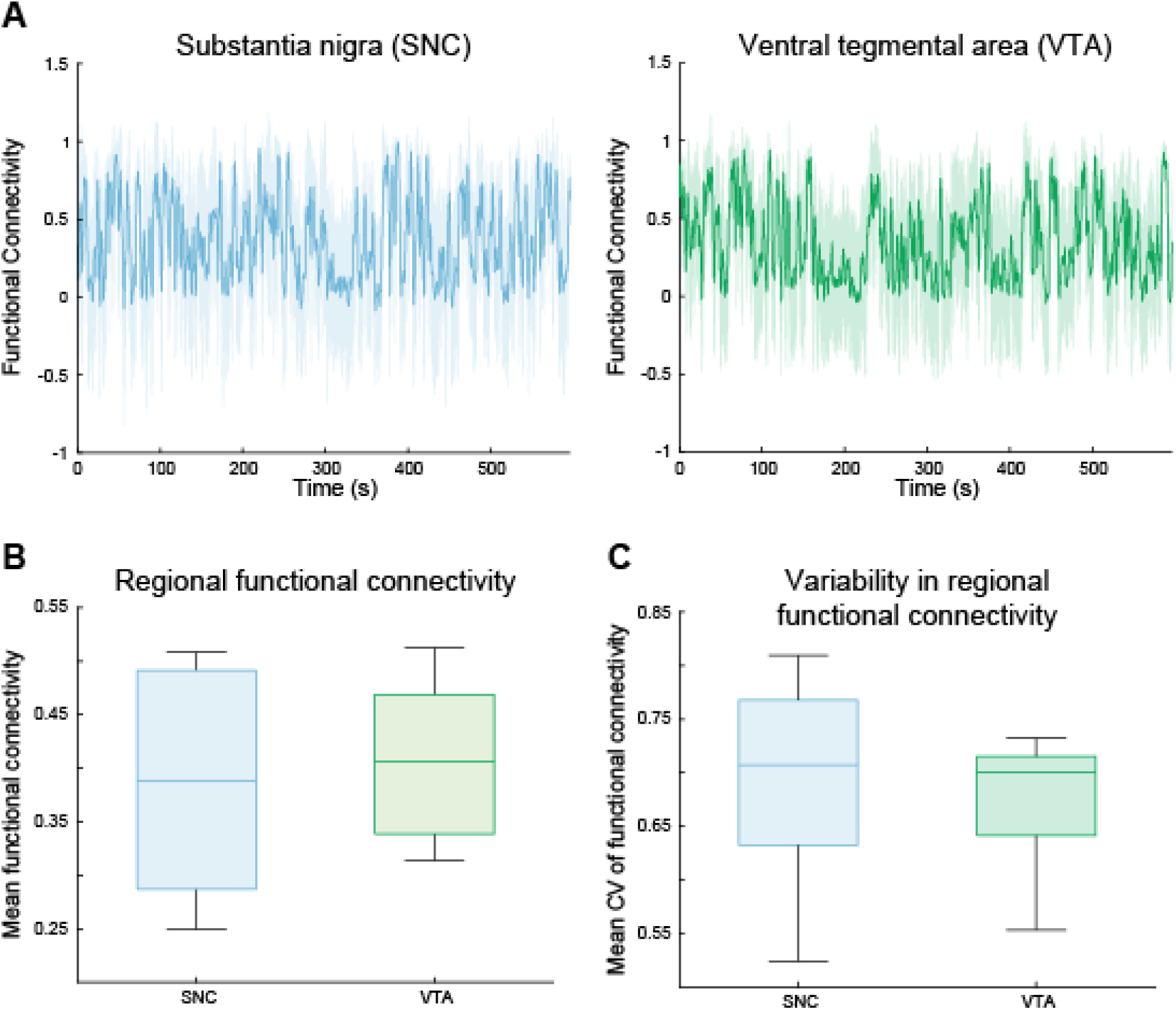
SNC and VTA exhibit similar patterns of dynamic functional connectivity. Intrinsic dynamic functional connectivity of dopamine neurons of the substantia nigra (SNC) and ventral tegmental area (VTA) exhibit similar patterns of functional connectivity. Functional connectivity is defined as the average pairwise Spearman’s correlation across all neuronal pairs. (**A**) Representative dynamic connectivity in the substantia nigra and ventral tegmental area. (**B**) Mean functional connectivity is similar in the substantia nigra and ventral tegmental area (mean FC = 0.386, 0.407 SNC, VTA respectively, two-tailed unpaired t-test, p = 0.7193). (**C**) Variability of functional connectivity (coefficient of variation, CV) is similar across the two regions (mean CV = 0.694,0.673 SNC, VTA respectively, two-tailed unpaired t-test, p = 0.6887) n = 5-8 sections from 5 animals.

We next asked whether these two regions would exhibit similar patterns in functional connectivity over time. To do so, we measured the coefficient of variation in the functional connectivity of each region and found similar patterns of functional connectivity within the SNC and VTA dopaminergic networks (Figure 2C, two-tailed unpaired t-test, p = 0.6887). Similar average and dynamic functional connectivity patterns in the VTA and SNC dopamine neurons suggest that each region has similar basal oscillatory processes, which have been theorized to underlie how dopaminergic networks encode information^8^. The limitation of average functional connectivity is that it does not provide information about the structure of the functional network, and specific properties these may confer.

In contrast to functional connectivity, complex network theory provides a well-accepted approach to investigate patterns of connectivity, which in turn facilitate or limit information transfer between dopamine neurons and across the network, that can reveal predisposition to failure - loss of information transfer within functional networks^41,100^. Therefore, we utilized the complex network analysis to examine the functional network properties of the SNC and VTA.

### Emergent variance of network hubs in the SNC and VTA

To investigate patterns of functional network connectivity in each region, we examined the strength of connections (magnitude of the correlated activity between neurons) in the SNC and the VTA. Within the observable neurons of the field of view in each region, we found a wide distribution of connection strengths between dopamine neurons, where some neurons would exhibit low connectivity to each neuron within the network, while others would be strongly interconnected, suggesting that hub neurons may exist within the dopaminergic neuronal network in the VTA and SNC (Figure 3A,B). Therefore, we calculated the nodal strength of each neuron by integrating the connection values to every other neuron per matrix and over time. Since more connections can arbitrarily inflate nodal strength values, temporal nodal strength means were normalized to the relative size of the network (Figure 3C). Similar to the pattern of network connectivity of the network visualizations in Figure 3A,B, we found differential patterns of connectivity between dopaminergic neurons of the SNC and VTA. In the SNC, we found skewing of the connection strength distribution to lower strength values. Whereas, VTA neurons exhibited stronger connections across the network compared to the SNC (mean normalized node strength (n) = 3.97 (89 neurons) - SNC, 5.24 (70 neurons) - VTA,two-tailed unpaired t-test, p = 1.89×10^−9^). Due to the insurmountable technical restraints using brain slices, one of the limitations of our approach is that we have computed node strength in relatively small neuronal networks- in each brain structure- comprising 10-20 nodes. Nevertheless, even in this limited scale our analyses revealed the SNC to be less densely interconnected than the VTA suggesting a potentially increased vulnerability of the region to insults.

**Fig. 3:**
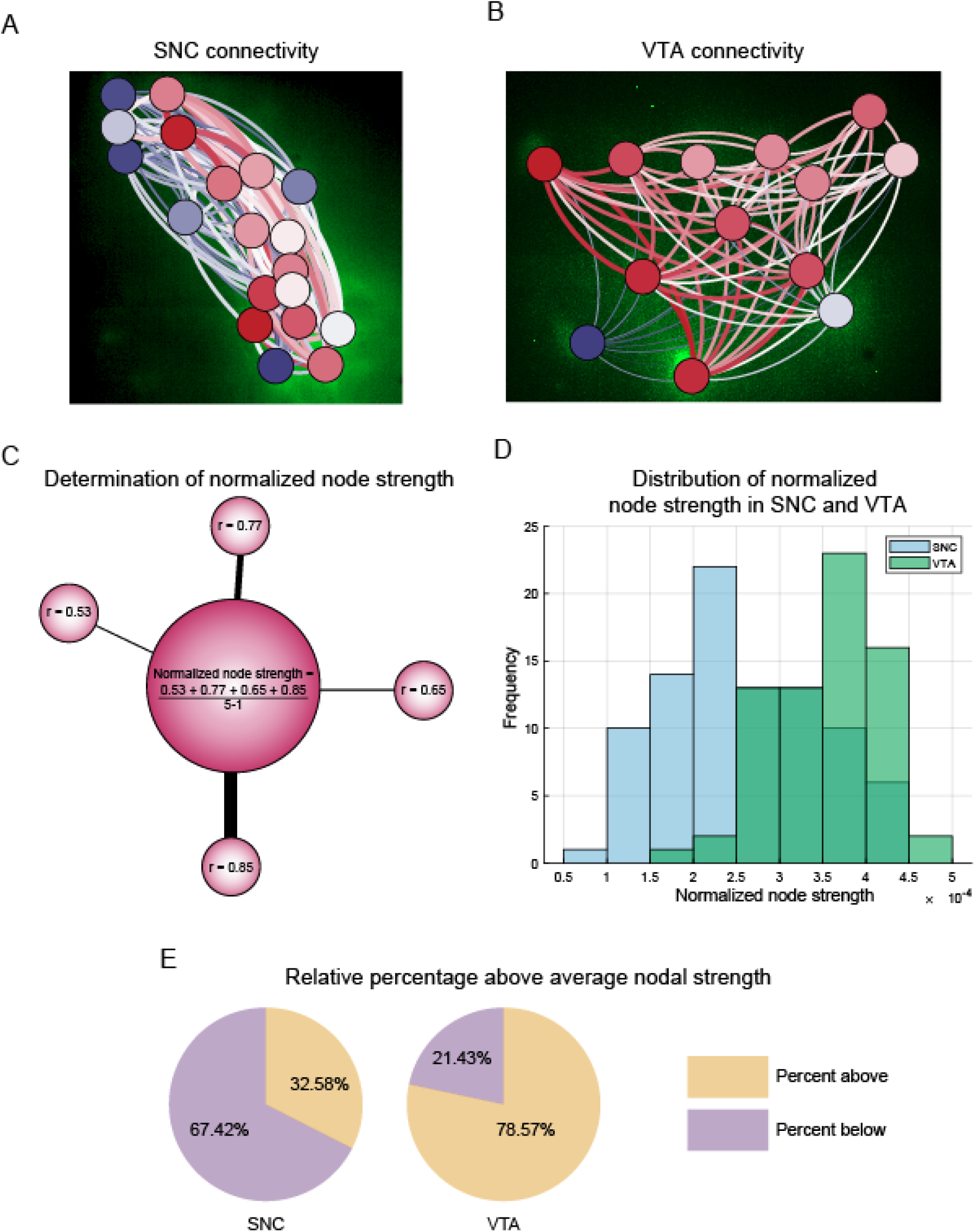
VTA dopaminergic networks are more strongly interconnected than the SNC. (**A,B**) XY-projection network graph of the functional connectivity matrices overlaid on SNC and VTA suggest differences in connection strength distributions. Connection strength, a neuron’s relative contribution to the network, is displayed through coloration (blue = minimum, white = mean, red = high). Neurons are overlaid with circles representing their weighted degree (node strength, blue = minimum, white = mean, red = high). Scales reflect distribution within the investigated network. (**C**) Normalized node strength is determined through integrating the magnitude of the correlation values of each neuron to all other neurons within the network, and then normalized by dividing by N-1, where N = total number of neurons within the network. (**D**) VTA neurons tend to exhibit larger normalized node strength values with a right skewed distribution while SNC exhibit a more Gaussian-like distribution centered on a lower value (mean normalized node strength (n) = 3.97 (89 neurons) - SNC, 5.24 (70 neurons) - VTA, two-tailed unpaired t-test, p = 1.89×10^−9^) Node strengths are pooled for each region from 5-8 slices from 5 animals. (**E**) Normalized node strength values were pooled for midbrain average node strength and then neurons were classified whether they exceeded or fell below this value. The VTA exhibits a significantly greater proportion of neurons exceeding this average midbrain node strength than SNC (78.57% - VTA, 32.58% - SNC, two-tailed unpaired t-test, p = 3.80*10^−13^).

The variances in nodal strength distributions between regions suggest that some neurons behave as network hubs, or neurons that significantly contribute more to the network than others. As nodal strength belongs to a group of network measurements known as nodal centrality, nodal strength provides a simplistic evaluation for hub-like properties of neurons within the region. We combined the weighted connection distributions (magnitude of Spearman’s rho between each neuron) of both regions to determine an average nodal strength value regardless of region, and calculated the relative percentages above and below the value for the SNC and VTA. Only a relatively small fraction of SNC neurons met this criterion (32.58%) while the vast majority of neurons in the VTA (78.57%) exceeded it. These results suggest that loss of hub neurons in the SNC may have more drastic consequences to network properties and its potential lower resilience to failure than a similar loss in the VTA, potentially underlying the differential sensitivity ascribed to the two regions^101,102^.

### Network properties of the SNC and VTA

Similar to other brain networks^103^, it is likely that dopaminergic network connectivity is non-random in functional structure. However, functional network topology can emerge from randomly interacting neurons^104,105^. Therefore, we sought to determine first whether real network properties of the SNC and VTA are distinct from chance events. To determine whether the dynamic properties of each region deviate from a random network of equivalent size, each property was evaluated against a random weighted network (or random lattice, where appropriate) of equivalent size and functional connectivity magnitude. Importantly, these random networks were generated through either a rewiring parameter performed for 1000 iterations randomly or with latticization^106,107^. Dopaminergic networks of the SNC and VTA were then assessed for network assortativity (preferential connectivity), modularity (delineation of clear non overlapping groups), path length (average network distance between nodes), and clustering coefficient (higher order interactions and functional segregation).

We first assayed network assortativity, which ascertains preferential attachment of a neuron to other neurons within the network. Specifically, if a neuron with many connections is primarily is connected to other neurons with many connections (preferred connectivity to neurons with similar connectivity), that neuron will have a positive assortativity value, whereas if a neuron with many connections primarily connects to neurons with only one connection (preferred connectivity to neurons with dissimilar connectivity), that neuron will have a negativity assortativity value. Importantly, positively assortativity has previously been shown to provide robustness and resilience to networks^108,109^ while negativity assortativity increases speed of information transfer^110,111^. Therefore, examination of network assortativity provides critical information about resiliency and speed of information transfer in the VTA and SNC dopaminergic neurons/network. We found that the SNC and VTA dopaminergic networks both exhibit weakly positive assortativity (random networks generally exhibited negative assortativity), suggesting that neurons preferentially connect to neurons with similar connectivity properties that oscillate between positive and negative values over time (Figure 4A-C).

**Fig 4:**
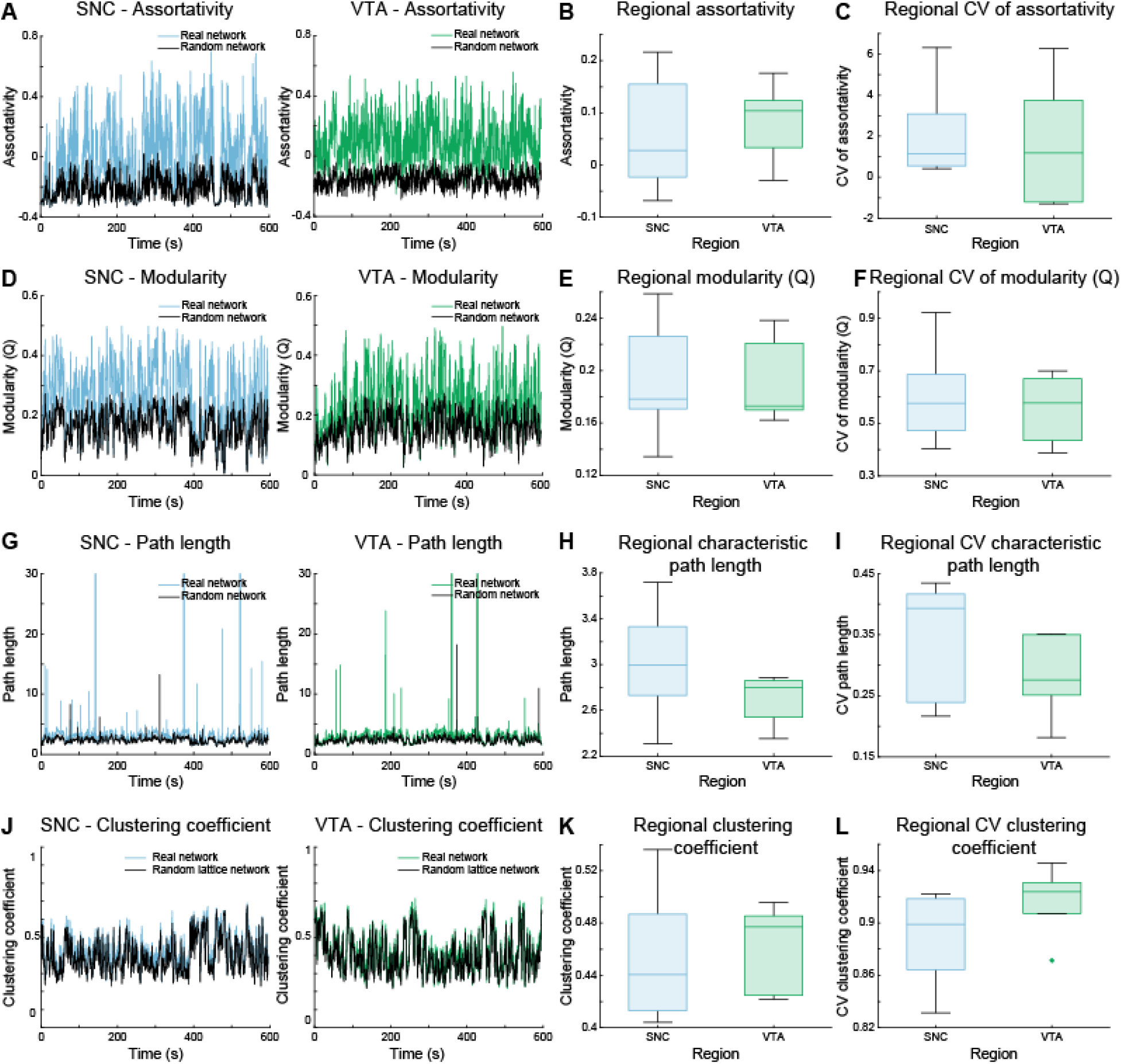
Complex network analysis reveals non-random ordered topology of SNC and VTA dopaminergic networks. We investigated regional dynamic network properties to determine whether they differ from those of a random network of equivalent size and connection distribution. Networks were assayed for (**A**) assortativity - preferential connectivity of a neuron to other neurons with similar (positive) or dissimilar (negative) number of connections, (**D**) modularity - the degree to which neurons form sub-specialized groups, (**G**) path length - the average number of connections separating any two neurons within the network, and (**J**) clustering coefficient - an triplet interactions between 3 neurons. Across each measurement, SNC and VTA networks exhibited similar network assortativity (**B**), modularity (**E**), characteristic path length (**H**), and clustering coefficient (**K**) that were significant from their randomized counterparts and exhibited similar dynamics (**C, F, I, L**). n = 5-8 sections.

We next quantified modularity of the network, or the degree to which neurons form specific groups that are more interconnected with each other than neurons outside the group, ranging from 0 to 1 where 0 represents no modular separation and 1 indicates a strong modular structure. Highly modular networks enable sub-specialization of the neurons within a network^112,113^. Similarly, both regions exhibited equivalent non-random consistent modular functional structures (Figure 4D-F), suggesting that modular segregation, and thereby sub-specialization of neuronal groups, occurs consistently in the SNC and VTA. To investigate the relationship of robustness and efficiency of information transfer, we evaluated the network beyond pairwise comparisons to triplets, determined by clustering coefficients - which assays the connectivity between 3 neurons simultaneously^65,114,115^. High clustering coefficients are associated with network robustness^116^. We additionally quantified global efficiency using path length, where a low path length suggests that information can quickly move between clusters facilitating temporally efficient transfer^116^. Path length and clustering coefficients in the SNC and VTA dopaminergic networks were significantly different from random networks and random lattices of equivalent size and connection distribution. Although significantly different, these followed patterns found in random networks, suggesting that more complex topology may be present, such as small-worldness^117,118^.

We next sought to investigate SNC and VTA dopaminergic networks for small-world topology, which is characterized by high clustering, similar to random lattice clustering, and low path length, similar to random path length^117,119^. Importantly, small-world networks can confer processing efficiency, resiliency and synchronization^44,101,117,120,121^. SNC and VTA networks met the standard criteria of a small-world coefficient (ω) between −0.5 and 0.5^119^ suggesting that both regions exhibit this property robustly across time (Figure 5A-C).

**Fig 5:**
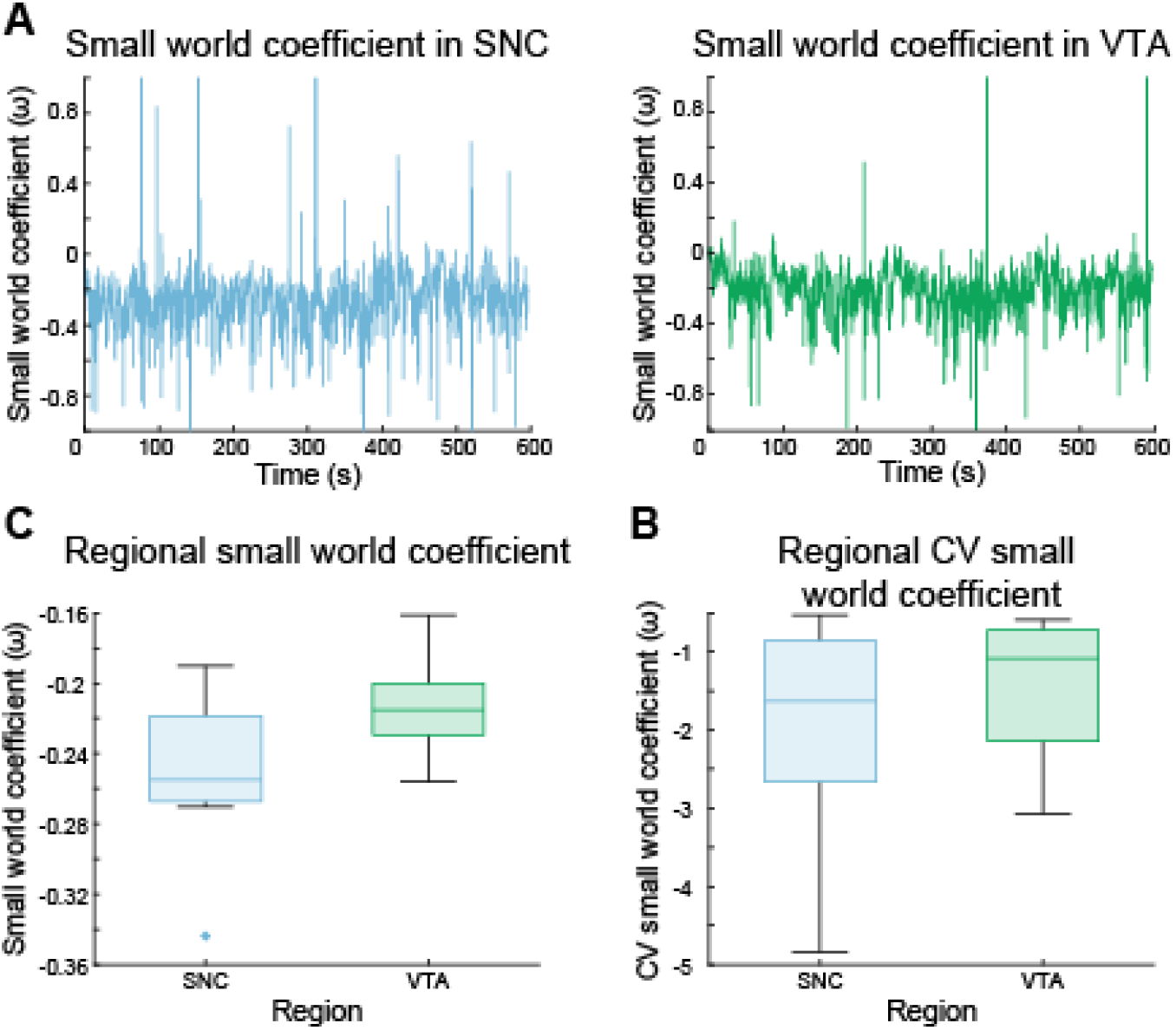
Dopaminergic networks exhibit small-world properties. SNC and VTA networks were assayed for small-world coefficients, characterized by high clustering (relative to a random lattice network) and low path length (relative to random network) over time. (**A**) Both SNC and VTA networks oscillate between similar lattice-like and random-like small-world topology. (**B,C**) SNC and VTA networks primarily exhibit lattice-like topology that is similarly stable over time (SNC versus VTA, small-world coefficient p = 0.1019, CV p = 0.4681, two-tailed unpaired t-test).

### SNC fragility and VTA resiliency in neuronal loss

Although the SNC and VTA exhibit surprisingly similar and robust dynamic network properties, the literature suggests these regions to be differentially sensitive to neuronal loss^30,50,56^. Furthermore, it remains unexplored whether loss of a single neuron significantly alters network properties. Therefore, we sought to investigate to what degree the functional network in each region is affected by the loss of a single neuron as determined by restructuring of network properties. To do so, we utilized simultaneous whole-cell patch clamp electrophysiology and calcium imaging to selectively hyperpolarize individual dopamine neurons to remove their contribution to the functional network activity (Figure 6A). Network measurements were performed at two stages: 1) at baseline firing activity, while the neuron is patched, but no current is injected (Fig 6A - note there is no change in the spontaneous firing of the patched neuron) and 2) during hyperpolarization, where a hyperpolarizing negative current is applied to suppress spontaneous firing activity (Fig 6A, note the absence of firing activity during the hyperpolarization step).

**Fig. 6.**
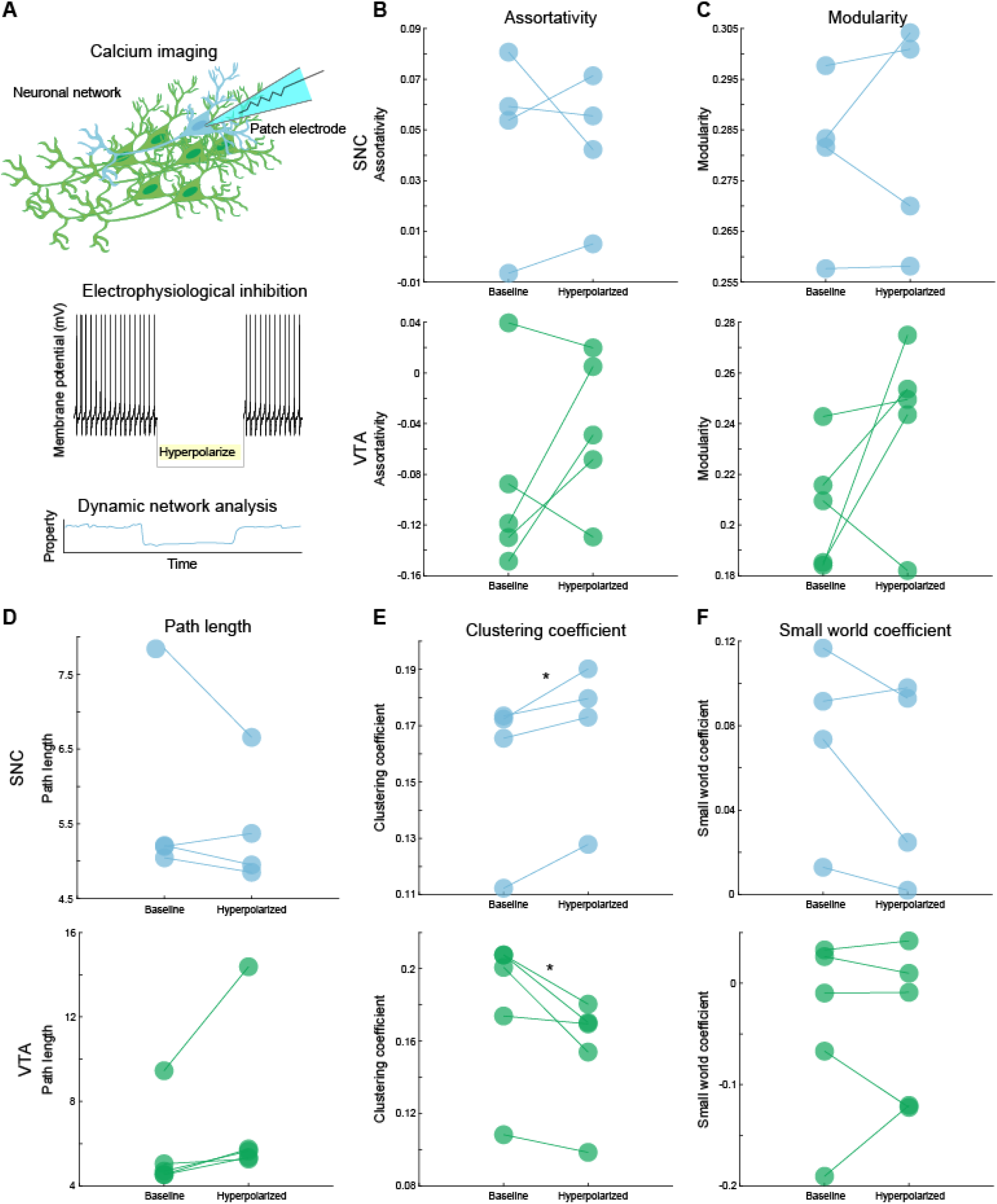
Suppression of individual neuronal activity increases network clustering in SNC but decreases network clustering in the VTA. (**A**) Top - Individual neurons within the field of view are patched sequentially with simultaneous calcium imaging. Middle - During the imaging paradigm, neurons are hyperpolarized to suppress their activity and eliminate contribution to the network. Bottom - networks are analyzed with the neuron in the network (baseline) and while activity is suppressed (hyperpolarized). The hyperpolarized neurons were excluded from network measurements. Network assortativity (**B**), modularity (**C**), path length (**D**), and small-world coefficient (**F**) remained stable across the SNC and VTA when individual neurons were hyperpolarized (hyperpolarized versus baseline, assortativity, p = 0.8127 SNC, p = 0.2418 VTA, modularity, p = 0.6604 SNC, p = 0.1792 VTA, path length, p =0.2947 SNC, p = 0.1249 VTA, small-world coefficient, p = 0.3264 SNC, p = 0.5243 VTA, paired samples two tailed t-test). Clustering coefficient (**E**) significantly differed in both regions (hyperpolarized versus baseline, p = 0.0271 SNC, 0.0353 VTA, two-tailed paired samples t-test), but were opposite in valence. n = 4-5 networks per group, * p < 0.05.

Intriguingly, the networks retained many of their properties such as assortativity, which ascertains whether neurons preferentially connect to neurons with similar connection distributions, and modularity, which measures the degree to which the network forms sub-groups (Assortativity, p = 0.8127 SNC, p = 0.2418 VTA, modularity, p = 0.6604 SNC, p = 0.1792 VTA). Strikingly, we found that loss of an individual neuron in SNC increases clustering, suggesting that SNC neurons become more coordinated following neuronal loss to potentially overcome the loss and likely increases metabolic burden. Conversely, VTA networks exhibited significantly decreased clustering coefficients, which may provide resiliency to the region by preventing failures from cascading across the network.

### Dopamine transporter modulation regulates dopaminergic network properties

Dopaminergic signaling is tightly regulated through dopamine transporter mediated reuptake. Previously we and others have shown that methamphetamine increases the DAT-dependent sodium current, intracellular calcium levels leading to increased firing frequency of dopamine neurons^61,62,122–125^. These data led us to the hypothesis that regulation of network properties may be under direct control of dopamine transporter activity. In order to test this hypothesis, we used methamphetamine to stimulate dopamine transporter activity. To determine specificity, these experiments were performed in the presence or absence of an antagonist cocktail that blocks the synaptic receptors D1, D2, GABA_A_, GABA_B_, AMPA, and NMDA, as well as a cocktail supplemented with a dopamine transporter inhibitor. This experimental paradigm allows for the delineation of which properties depend directly on dopamine transporter specific mechanisms, receptor mediated, and those which can occur independently. We found methamphetamine increased assortativity in both methamphetamine alone (Figure 7A, drug versus baseline, p = 0.0152, paired samples two-tailed t-test) and with antagonists (drug versus baseline, p = 0.0361, paired samples two-tailed t-test), but not in the presence of nomifensine blockade of dopamine transporter (drug versus baseline, p = 0.0503, paired samples two-tailed t-test), although a similar trend is observed. A similar effect was observed for small-world coefficients (Figure 3E, drug versus baseline, methamphetamine alone p = 0.0125, methamphetamine with antagonist cocktail p = 0.0046, methamphetamine, nomifensine, and antagonists p = 0.0713, paired samples two-tailed t-test). Since methamphetamine decreases dopamine uptake^126,127^, increases dopamine efflux^60,61,128^, and increases firing activity of dopamine neurons^62,89,129^, it is not surprising that methamphetamine exposure increased modularity and reduced network clustering across all conditions (Figure 7B,C drug versus baseline, modularity methamphetamine alone p = 0.0104, methamphetamine with antagonist cocktail p = 0.0012, methamphetamine, nomifensine, and antagonists p = 0.0323, clustering coefficient - methamphetamine alone p = 0.0159, methamphetamine with antagonist cocktail p = 0.0068, methamphetamine, nomifensine, and antagonists p = 0.0488, paired samples two-tailed t-test), while path length remained consistent throughout (Figure 7D, methamphetamine alone p = 0.9238, methamphetamine with antagonist cocktail p = 0.3215, methamphetamine, nomifensine, and antagonists p = 0.1489, paired samples two-tailed t-test). These results collectively suggest that modulation of dopamine transporter activity can regulate dopaminergic network properties.

**Fig. 7.**
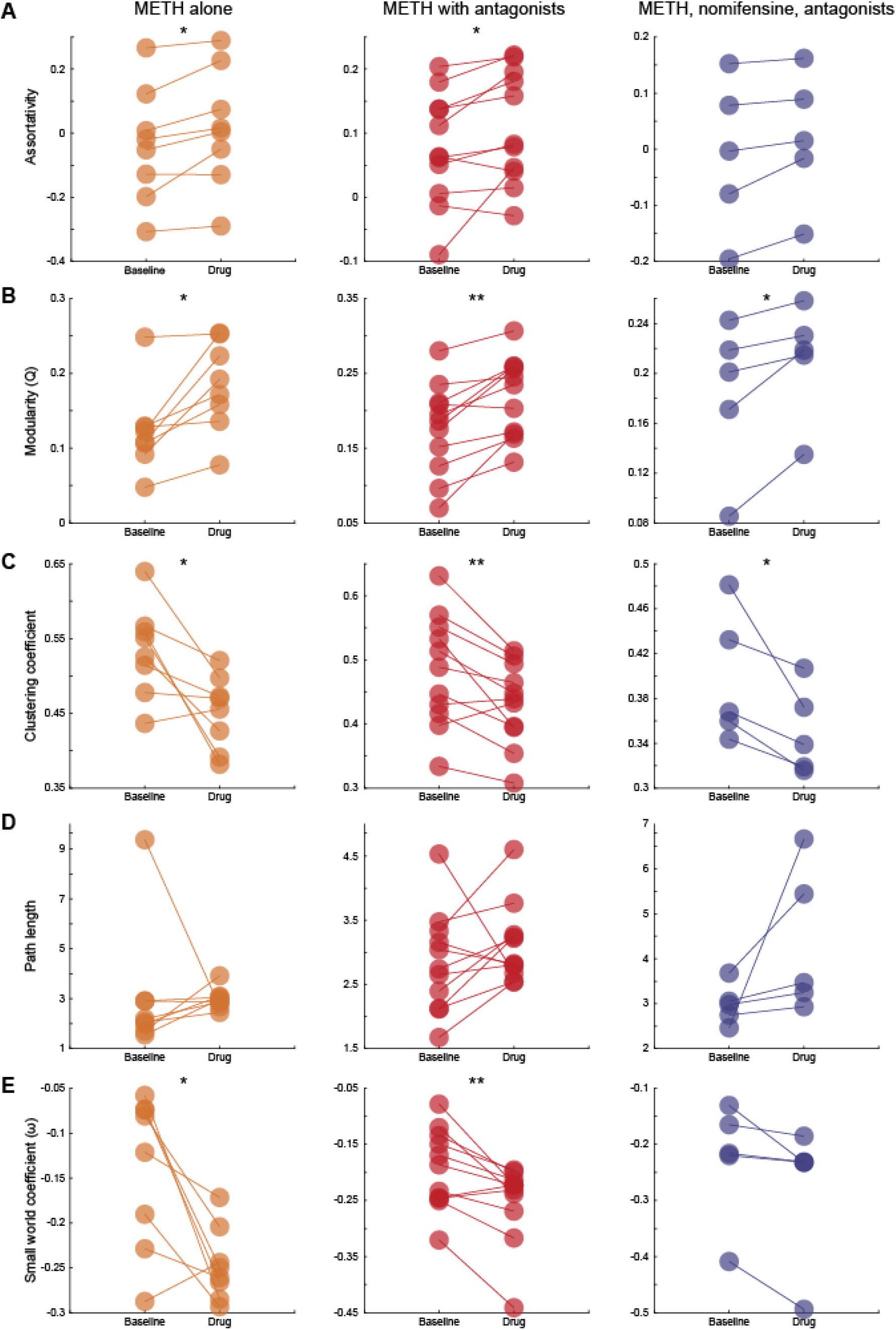
Dopamine transporter modulation regulates dopaminergic network properties. (**A**) Dopaminergic networks stimulated with methamphetamine exhibit increased assortativity whether methamphetamine alone or with antagonist cocktail blocking : D1, D2, GABA_A_, GABA_B_, NMDA, AMPA receptors) is applied (methamphetamine alone p = 0.0152, methamphetamine with antagonists p = 0.0361, p = 0.0503 methamphetamine with nomifensine and antagonists, two-tailed paired samples t-test). (**B**) Network modularity increased across all conditions (methamphetamine alone p = 0.0104, methamphetamine with antagonists p = 0.0012, methamphetamine with nomifensine and antagonists p = 0.0323, two-tailed paired samples t-test). (**C**) Clustering coefficients of dopaminergic networks significantly decreased across all conditions (methamphetamine alone p = 0.0159, methamphetamine with antagonists p = 0.0068, methamphetamine with nomifensine and antagonists = 0.0488, two-tailed paired samples t-test). (**D**) Path length remained the same across all stimulation conditions (methamphetamine alone p = 0.9238, methamphetamine with antagonists p = 0.3215, methamphetamine with nomifensine and antagonists p = 0.1489, two-tailed paired samples t-test). (**E**) Methamphetamine stimulation promoted lattice-like small-worlds in dopaminergic networks, but not when dopamine transporter is occluded (methamphetamine alone p = 0.0125, methamphetamine with antagonists p = 0.0046, methamphetamine with nomifensine and antagonists p = 0.0713, two-tailed paired samples t-test). n = 5-11 networks per group, *p < 0.05, **p < 0.01.

## Discussion

Dopaminergic neurons and by proxy their regional networks exhibit a rich diversity of neurochemical and electrophysiological properties^3,4^. The focus of the field often relies upon results from individually recorded neurons^4,35,59,62,98^, or distinctions derived from immunohistochemistry^3^ or transcriptomics^1,2^ that cannot capture whether the activity in one neuron influence the activity of local neurons and ultimately the output of dopaminergic network. To address these limitations, we utilized simultaneous highspeed calcium imaging and electrophysiology to isolate interregional differences in interactions of spontaneously active dopaminergic neurons of neighboring dopaminergic nuclei, the SNC and VTA at baseline and following exposure to methamphetamine known to increase dopamine transmission via multiple mechanisms.

Our results show for the first time that the dopaminergic networks of the substantia nigra (SNC) and ventral tegmental area (VTA) exhibit stable and rich dynamic network properties. We identified increased prevalence of network hubs within the VTA compared to the SNC, differential responses to loss of a single neuron, and modulation through dopamine transporter dependent and independent mechanisms.

Functional networks emerge from local interactions between neurons^46,47,49^. In dopaminergic networks, dopamine neurons have the machinery to directly interact through both synaptic and non-synaptic signaling^65,91,130^. However these studies do not account for dopaminergic network properties and dynamics, and how they differ by brain region. Identification of regional differences offer greater understanding of differential sensitivity of dopaminergic regions, such as loss of SNC dopamine neurons in Parkinson’s disease^30,55,56^. Within the subsets of each regional area sampled, we found an overrepresentation of hub-like neurons in VTA dopaminergic networks than in the adjacent SNC. Therefore, the event of loss of a hub neuron in the SNC would have a greater detriment to behavioral outputs that are circumscribed to the SNC. Finally, although the observed regions represent a small fraction of the total area of the region, which is the general limitation of live cell imaging in the randomly selected regions in the SN and VTA, our computational analyses still revealed non-trivial differences in network topology between regions. Future work will determine the degree to which this is consistent across the three-dimensional axes (rostral-caudal/medial-lateral) of the SNC and VTA.

The observed differences in basal network properties led us to test the hypothesis that neuronal loss should differentially affect these two regions. Selective removal of neurons from functional networks through electrophysiological suppression resulted in surprisingly increased clustering of the SNC with decreased clustering in the VTA (Figure 6E). While high clustering, a higher order measure of connectivity^114,115^, is associated with network robustness^116^, increased in clustering suggest that SNC networks respond to neuronal loss by increasing functional segregation^100^ to potentially maintain network output, while the VTA responds by becoming more dispersive. Increased clustering between neurons in the SNC suggest an increase of neuronal resources that would compensate the contrasting decreases in average firing rates in previous literature^131^, creating metabolic stressors that may predispose further neuronal dysfunction and consequently network failure^50,132,133^. Thus, the network attempting to maintain synchrony by temporally aligning may then contribute to dysfunction of other nuclei within the basal ganglia^134,135^. Conversely, the response found within the VTA would suggest robustness resilience to cascades of energy usage or pathogens^100,136^. However, loss or dysfunction of VTA dopamine neurons also underlies some neuropsychiatric dysfunction^137^. Therefore, given enough dispersion of VTA dopamine networks, the outcome of loss of higher order synchrony and functional specialization may underlie these symptoms.

Beyond single neurons, dopaminergic networks feature volume transmission and therefore can interact with both proximal and distal targets^75,82^. Dopamine transporter regulates the degree to which dopamine diffusion can occur through the reuptake of dopamine and ending dopamine signaling^77,125^. Furthermore, influx of dopamine or methamphetamine through the transporter produces depolarizing currents that alter the firing properties of these neurons^59,62,91^, which both transporter activity and firing rate are critical to dopaminergic synchronization^8,87^. Disruption of dopamine transporter function would therefore likely cascade across the network, disrupting network topology. Our data support the interpretation that network structure is under tight control by dopamine transporter, as all but one metric (path length) significantly altered through manipulation of dopamine transporter, and these effects persisted under blockade of many of the receptor types found within the ventral midbrain, but not when the transporter was blocked (Figure 7). Therefore, dopamine transporter can act as a master regulator of dopaminergic networks through regulation of dopaminergic signalling. Additionally, this provides a direct pharmacological target that may be able to revert dopaminergic networks with dysfunctional properties, such as those found in Parkinson’s disease^138^ and addiction^139–141^. However, as the experiments were performed in a cocktail of antagonists, future experiments will be necessary to provide more nuanced interpretations to the regulation of dopaminergic networks through simultaneous polypharmacy approaches of multiple targets that have been shown to alter network synchrony^142^. Importantly, while acute midbrain slices provide regional investigations into the regulatory mechanisms within each region, they do not account for active afferent input during behavioral state-dependent changes observed in freely behaving animals^143–145^. Therefore, future experiments will require in-vivo monitoring of network activity to determine whether and how dopaminergic networks of the SNC and VTA shift their topology to produce or respond to state-dependent changes.

These results expand and complement the current body of literature from the firing rates of individual neurons and synchrony to networks of interacting dopaminergic neurons whose remarkable heterogeneity give rise to exquisite regional differences, both in sensitivity and resiliency. Therefore, our approach provides a foundation for future experiments into the functional structure of dopaminergic networks and how changes in functional structure underlie behavior and disease.

## Materials and Methods

### Animals

DAT^IRES*cre*^ and Ai95(RCL-GCaMP6f)-D knock-in mice were obtained from The Jackson Laboratory (Stock number: 006660 (DAT^IRES*cre*^), 024105 (Ai95D), Bar Harbor, ME, USA). Only animals exhibiting bright EGFP fluorescence were used for experiments. Approximately equal numbers of males and females were used for each experimental condition. All animals were maintained in the University of Florida animal facilities. All experiments were approved by the Institutional Animal Care and Use Committee at University of Florida. Animals were housed under standard conditions of 22-24°C, 50-60% relative humidity, and a 12 hour light/dark cycle.

### Preparation of acute midbrain slices in imaging

Postnatal day 28-42 DAT-cre/loxP-GCaMP6f mice of either sex were used. Mice were decapitated and brain extracted immediately into ice-cold oxygenated ACSF containing (in mM) 126 NaCl, 2.5 KCl, 2 CaCl_2_, 26 NaHCO_3_, 1.25 NaH_2_PO_4_, 2 MgSO_4_, and 10 dextrose, equilibrated with 95% O_2_–5% CO_2_, 300-310 mOsm, and pH adjusted to 7.35-7.4. Brain was cut rostral to PFC in the coronal plane to create a flat surface to glue to the cutting stage. Brain was then submersed in fresh ice-cold ACSF as previously described. 200-300 uM sections containing either the substantia nigra or ventral tegmental area were cut using a DSK MicroSlicer Zero 1N and transferred to a recovery chamber at 32-34°C and allowed to recover for 45-60 minutes and maintained at room temperature (22-24°C) until used.

### Electrophysiological recordings and analysis

To facilitate electrophysiological recordings, the slicing preparation was modified with slicing performed in a sucrose-sodium-replacement solution containing (in mM) 250 sucrose, 26 NaHCO_3_, 2 KCl, 1.2 NaH_2_PO_4_, 11 dextrose, 7 MgCl_2_, 0.5 CaCl_2_ and recovery (ACSF) was supplemented with 10 μM MK-801 to reduce glutamate toxicity. Slices were transferred to a low profile open diamond bath imaging chamber (RC-26GLP, Warner Instruments, Hamden, CT, USA) under constant perfusion of ACSF (described previously) at 36-37°C at a rate of 2 ml/minute. Dopamine neuron cell bodies were identified first by GCaMP6f fluorescence through a 40x water immersion objective on an Eclipse FN1 (Nikon Instruments, Inc., Melville, NY, USA) upright microscope using a Spectra X light engine (Lumencor, Beaverton, OR, USA) equipped with a 470/24 nm solid-state illumination source and recorded using a 12-bit Zyla 4.2 sCMOS camera (Andor Technology, Ltd., Belfast, Northern Ireland). Neuronal morphology was then visualized using infrared differential interference contrast (IR-DIC). Borosilicate glass capillaries (1.5 mm O.D. Sutter Instrument Company, Novato, CA, USA) were pulled on a P-2000 laser puller (Sutter Instrument Company, Novato, CA, USA) to an open tip resistance of 4–10 MΩ and filled with a potassium gluconate-based internal solution comprised (in mM) 120 K-gluconate, 20 KCl, 2 MgCl_2_, 10 HEPES, 0.1 EGTA, 2 Na2ATP, 0.25 NaGTP, 5 biocytin, 290-295 mOsm, pH adjusted to 7.25-7.30. Recordings were made using an Axon Axopatch 200-B microelectrode amplifier and digitized with a Digidata 1440a through ClampEx 10.2 software (all Molecular Devices, LLC, San Jose, CA, USA). Analysis of electrophysiological recordings was performed with custom code written for this project using Python 3.7 and the pyABF module^146^.

### Calcium imaging

Acute coronal slices were illuminated as described previously (see *Electrophysiological recordings* section). Images captured at 20-25 frames per second with no delay between frames. Videos were acquired either simultaneously or independent of electrophysiological recordings, depending on the experiment performed.

### Image correction, segmentation and signal preprocessing

Nikon nd2 files were imported into ImageJ2 via the Bio-Formats importer. Motion correction was performed as necessary with the moco plugin^147^. Cell bodies of spontaneously active dopamine neurons were manually segmented and exported to CSV. CSV files were imported into Matlab and signals were smoothed using the loess method (span = 0.5% of data points). Signals were normalized using relative fluorescence by the following calculation:

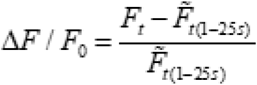

Where F at time *t* is normalized to the baseline measurement. The baseline is calculated as the median value of the first 25 seconds of the recording on a per neuron basis.

### Dynamic functional connectivity and network analysis

Dynamic functional connectivity was computed using pairwise Spearman’s correlation coefficients (MATLAB function: corr, type: Spearman) calculated over a sliding 100 frame window with a 1 frame step size between neurons. Since no threshold is privileged and arbitrary thresholds affect network measurements^94,96,148^, multiple proportional thresholds were applied to each correlation matrix from 5% to 100% at 1% intervals. Thresholds were excluded if all connections remaining were to one neuron or if less than 4 neurons remained in the network. For modularity and path length measurements, negative correlations were set to 0 to enable proper computation. All network analysis utilized the weighted function versions from the Brain Connectivity Toolbox^100^. Control weighted networks were created by randomizing the connection topology while maintaining the size, density, and degree distribution^106^, or latticized^107^ where appropriate. Network metrics were calculated at each threshold for both real and randomized networks and then averaged.

For all networks, usage of node refers to a neuron. Normalized network strength was defined as the sum of all weights per neuron divided by the number of neurons in the observed network^100^ as defined by:

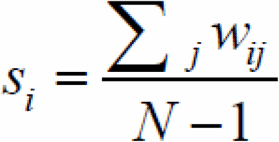

Importantly, normalized network strength was calculated using the full correlation matrix without thresholding.

Network assortativity, an evaluative measurement of preferential connectivity to neurons with similar or dissimilar connection strength^65,108,149,150^. Assortativity (r) was calculated as defined by:

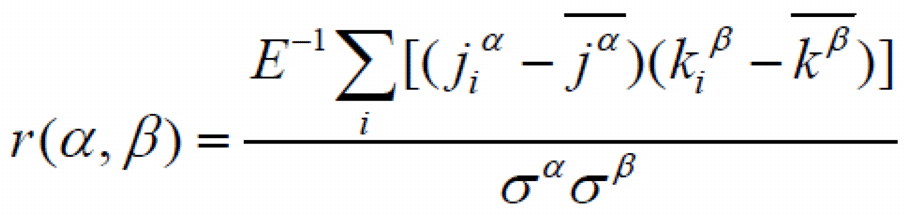

Where the nodal strength of neuron jα (source) and nodal strength of neuron jβ (target) at connection i of all connections (E). With σ defined similarly for α and β be:

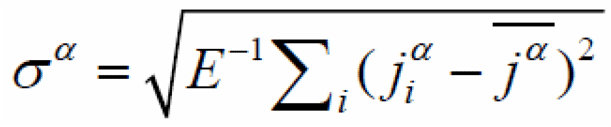

Network clustering, an extension on pairwise-correlation to triplet correlations^65,114^, was determined by calculating clustering coefficients^151^. Clustering coefficients were calculated as defined by:

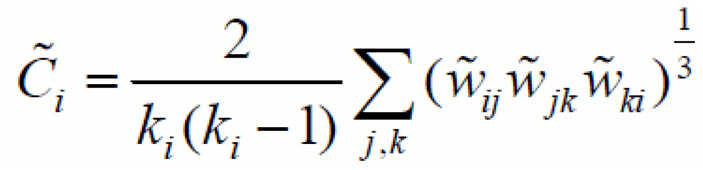

Where k_i_ is the degree of node i and the weight w between nodes i and j, computed across the triangle inclusion of nodes j,k.

Characteristic path lengths, the average minimum number of connections between any two neurons in the network that must be traversed to connect them^100^, infinite path lengths were omitted from the calculations. Following the original definition by of a small-world network^117,119^, the small-world parameter (ω) was calculated as defined by

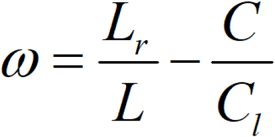

Where L and C refer to path length and clustering coefficient and Lr is the path length of a weighted random lattice and C_1_ is clustering of a weighted random network.

Network modularity, the existence of communities around neurons is quantified by the Q value index^152,153^.

### Statistical analysis and network graph construction

All statistical analysis was performed in MATLAB (version 2020a, Mathworks, Cambridge, MA, USA). Two-sample t-tests were used where appropriate. Effects were determined as significant at the alpha = 0.05 level. Undirected network connections were imported with corresponding XY positions from the image plane into Gephi v0.9.2^154^.

## Data and Code Availability

All raw ROI fluorescence data, and processed functional connectivity and network metrics are provided in MAT files (Matlab). Custom Matlab code is intended to run as compiled code on a high-performance computing cluster. Matlab livescript versions of the code, with raw and processed data, are available upon request.

## Acknowledgements

Douglas Miller is supported by T32-NS082128. Dr. Habibeh Khoshbouei is supported by R01-DA026947 and R01-NS071122. We would like to thank Dr. Jeff Beeler for his insightful comments that improved the quality of the manuscript.

